# dnaudit + pydnaweb: A lightweight text-based planning and documentation workflow for genetic cloning with automatic verification

**DOI:** 10.1101/2025.05.31.657172

**Authors:** Patrícia Ataíde, Faezeh Ghasemi, Paulo César-Silva, Sandra Paiva, Björn Johansson

**Affiliations:** Centre of Molecular and Environmental Biology, Department of Biology, University of Minho, Braga, Portugal; Institute of Science and Innovation for Bio-Sustainability (IB-S), University of Minho, Braga, Portugal

**Keywords:** Pydnaweb, dnaudit, Documentation, Pydna, Python, Molecular Biology

## Abstract

Life science research often depends on the construction and analysis of recombinant DNA molecules, where sequence accuracy is critical. However, the field continues to face a reproducibility crisis, partly due to the lack of comprehensive, systematic, and verifiable documentation of genetic constructions. Although most cloning procedures are deterministic and theoretically describable in a complete and unambiguous way, published methods are typically described in a form free narrative, making them laborious to reproduce and assess for completeness. Tools like the Python package Pydna support programmable and reproducible cloning strategies but require coding expertise, which can be a barrier for some users. To address this, we developed Pydnaweb and dnaudit, two open-source and complementary web tools that build on Pydna. Pydnaweb offers simulation of unit operations such as PCR and restriction digestion providing results in text format. These results can be collected and combined to form complex cloning strategies in a bottom-up approach. Dnaudit can verify such collections for internal consistency and that the cloning strategy meet a specific goal such as the expression of a protein sequence. The tools are design for a low barrier of entry, and they can be used separately. This workflow enables fully automated validation, providing a no-code, reproducible solution for documenting and sharing molecular cloning workflows. These tools ease compliance with FAIR principles and align with emerging standards for the transparent and reproducible sharing of scientific methods and data.

## 1. Background

The open availability of research results funded by public resources brings significant social and economic benefits. Funding entities are increasingly emphasizing creating data management plans, which are designed to assist and stimulate researchers to carefully consider how they document, manage, organize, and store their data throughout the research process (1,2).

In 2021, a survey was conducted to identify the factors contributing to poor reproducibility across various scientific disciplines. The primary issue identified was not the complexity of understanding laboratory records for most participants. Instead, the main contributors were found to be insufficient information in the materials and methods sections or experimental protocols, as well as a lack of comprehensive metadata regarding the experiments (3). These affected approximately 73-75 % of the participants (3), which highlights the critical importance of documentation in scientific research, particularly in reproducibility in molecular biology. Given the tangible nature of the objects and the inaccessibility to methodological details, accurately translating procedures and experimental outcomes in molecular biology becomes increasingly challenging (4).

Modern biological experiments often necessitate constructing new DNA molecules for various cellular purposes, such as protein expression. However, the planning stage of these constructs still heavily relies on using DNA sequence editors, such as SnapGene® (from Dotmatics; available at snapgene.com) (5-7) or the cloud-based platform Benchling (available at benchling.com) (8,9). Although these tools offer user-friendly features, their proprietary, licensed, and closed-source nature limits accessibility, transparency, and reproducibility, particularly in resource-limited environments or in the context of long-term projects. In such cases, the inherent complexity can increase the likelihood of errors, omissions, and a reliance on human interpretation when reconstructing cloning strategies. Notably, the lack of clarity and completeness of molecular cloning is especially concerning and ultimately avoidable, as DNA construction is, in principle, inherently deterministic.

This challenge is not merely theoretical. For instance, in a study conducted in 2021, the authors successfully reengineered the mevalonate (MVA) pathway in *Saccharomyces cerevisiae* to enhance squalene production. Briefly, expression vectors encoding key enzymes of the pathway were constructed using traditional restriction enzyme-based methods. Subsequently, guide RNA plasmids were assembled via In-Fusion cloning, and PCR-derived expression cassettes were integrated using CRISPR-mediated genome editing. Despite the clarity of the experimental rationale, essential details such as the restriction enzymes used for the gene cassette construction, plasmid maps, or complete sequence information were not provided. As a result, reproducing the DNA constructs would require either direct correspondence with the authors or speculative reverse engineering.

Towards addressing these challenges, various tools - both commercial and open-source - have been developed to optimize the design, assembly, and documentation of DNA constructs. The commercial tool j5 automates the design of complex DNA assemblies, optimizing for cost and efficiency, while simulating multiple assembly methods to select the most suitable option (10). Moreover, DNAda complements j5 by supporting high-throughput DNA assembly; it integrates with robotic systems for precise liquid handling and active sample tracking, thereby streamlining the entire process from in silico design to physical implementation (11). In contrast, open-source tools offer increased flexibility and adaptability for a wider range of users. For example, Teemi is a Python-based platform that combines literate programming and machine learning to integrate key stages of the design-build-test-learn cycle within a single workflow, thereby facilitating reproducibility and ease of use in microbial strain construction (12). Furthermore, Biolegato provides an object-oriented graphical user interface (GUI) that enables automation and customization of bioinformatics analyses through programmable workflows (13). Additionally, FastCloneAssist offers a user-friendly solution for designing primers used in restriction- and ligation-independent cloning strategies, eliminating the need for extensive coding expertise (14).

Whereas tools such as DNAda, which enable the execution of more tasks in less time and with reduced error rates, are primarily designed to meet the demands of industrial-scale and high-throughput applications (11,15,16), others, including Teemi, FastCloneAssist, and BioLegato, offer more adaptable solutions suited to academic and exploratory research settings (12-14). Notably, although many of them aim to produce shareable outputs, these are often structured primarily for machine execution rather than for human readability. This loss of human interpretability can compromise the transparency and auditability of molecular workflows, ultimately hindering reproducibility and collaborative exchange.

This study aims to present Pydnaweb, an open-source web-based interface built on the Pydna library, which significantly facilitates the simulation and transparent documentation of DNA cloning workflows by using a strategy description that is human-readable, while also being easy to parse by a computer. By streamlining the design and analysis of cloning pipelines, Pydnaweb can enhance the speed, accuracy, and reproducibility of individual steps of molecular biology strategies, further supporting researchers in their daily workflows. In addition, we also introduce dnaudit, another open-source tool that is well-suited for version control with Git and is capable of verifying the completeness and accuracy of planned cloning procedures. In this paper, we present two detailed examples illustrating how Pydnaweb and dnaudit can be effectively utilized to generate comprehensive and transparent documentation that adheres to FAIR (Findable, Accessible, Interoperable, and Reusable) principles (**Figure 1**), thereby promoting reproducibility and data sharing in molecular biology research (2,17).

**Figure 1:**
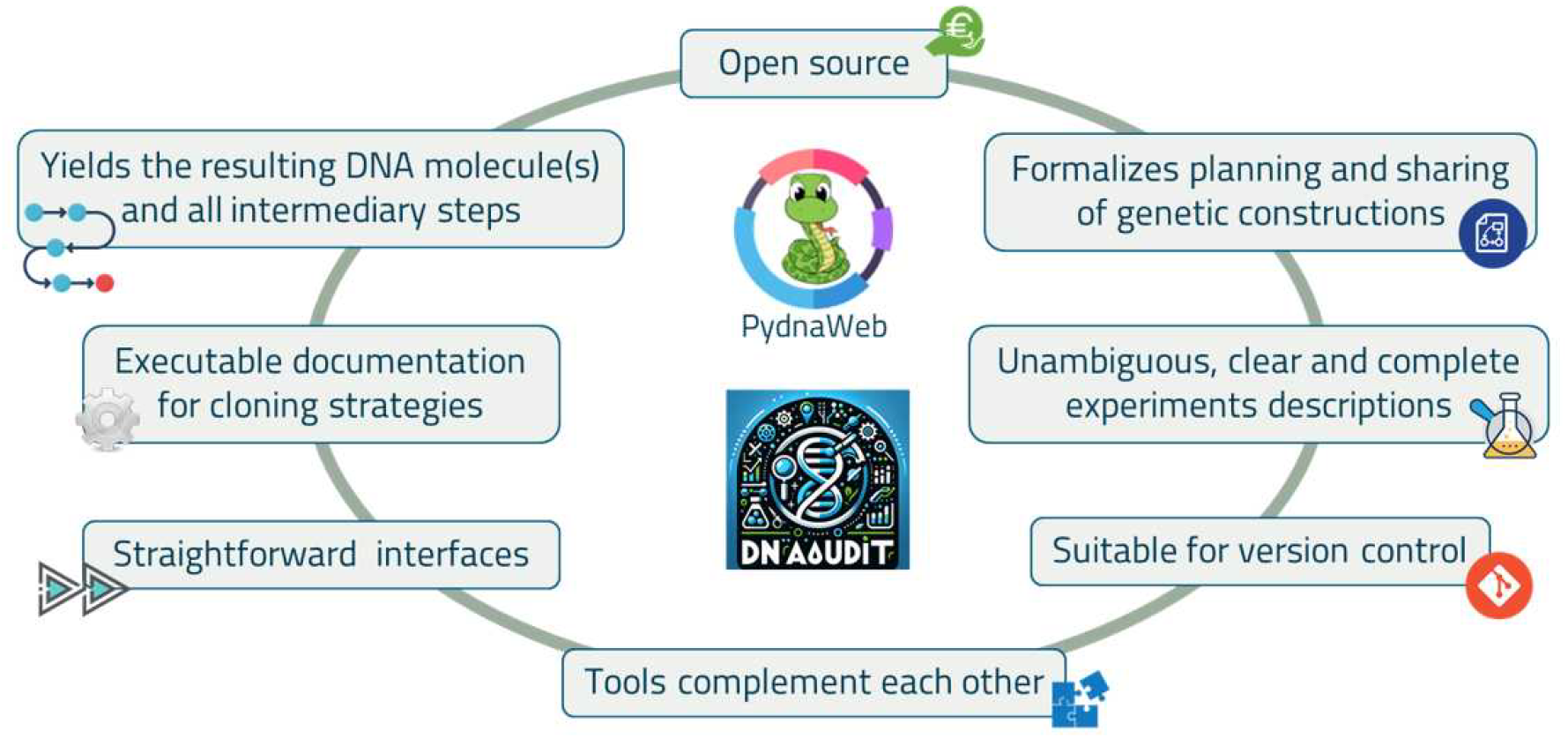
Key features provided by Pydnaweb and dnaudit, emphasizing their contributions to facilitating reproducible molecular biology workflows.

## 2. Results

### 2.1. Motivation for Pydnaweb and dnaudit

Biosciences are currently suffering from a reproducibility crisis that is exacerbated by the lack of systematic, complete, and verifiable cloning documentation. To address this issue, we evaluated existing tools (**Table 1**) to determine their suitability for our objectives: enabling reproducible molecular biology workflows and ensuring well-structured, comprehensive documentation. Achieving these goals would support more effective knowledge transfer within the research community and would foster trust in published results. Furthermore, improved reproducibility would facilitate collaboration by allowing researchers to build upon prior work, reducing redundant efforts and mitigating the risk of restarting projects due to irreproducible results.

**Table 1:** Comparison of molecular biology tools based on their support for structured documentation, reproducibility, and workflow automation. This table was compiled using available information from references 10-14.

| Tools | Components | User-Friendly | Workflow Automation | Visual Interface | Reproducibility Support | Structured Documentation |
| --- | --- | --- | --- | --- | --- | --- |
| j5 |  | ☑ | ☑ | ✗ | ⚠ (limited traceability) | ✗ |
| DNAda |  | ☑ | ⚠ (data-driven) | ✗ | ✗ | ✗ |
| BioLegato |  | ☑ | ⚠ (some scripting) | ☑ | ✗ | ✗ |
| Teemi |  | ☑ | ☑ | ☑ | ⚠ (limited output) | ☑ (Modular, machine-readable literate protocols) |
| FastCloneAssist |  | ☑ | ☑ | ☑ | ⚠ (not fully traceable) | ✗ |

Although these tools **(Table 1)** are user-friendly, accessible, and often provide helpful features such as visual design interfaces, automation of cloning steps, or integration with common lab workflows, none of them fully align with our objectives of generating clear, structured, and reproducible descriptions of molecular workflows. Nonetheless, the use of these tools can be valuable in many contexts, as their features are well-suited for specific tasks or workflows depending on the research needs. However, when it comes to reproducible and standardized documentation, most lack transparent outputs or the ability to trace and validate each step of a cloning strategy in a FAIR-compliant manner.

The Python package “Pydna” facilitates DNA manipulation procedures like primer design and DNA assembly simulation (18). While powerful, its reliance on coding poses a challenge for users who may not be proficient in programming. To overcome this limitation, most of the pydna Python package functionalities are exposed and available as a web service through a flask-based application, called Pydnaweb. Pydnaweb can simulate PCR, enzyme restriction, CRISPR digestion, sticky-end ligation, and homologous recombination, while facilitating standardization of protocols and cloning strategies.

When using Pydnaweb for a cloning simulation, a report in light plain text format is generated. This report provides a detailed description of the molecular biology unit operation that was simulated. The step-by-step process of generating a genetic construct can therefore be obtained, ensuring reproducibility, while also supporting the preservation, storage, and sharing of genetic material. Such a construct may include, for example, a plasmid assembled from fragments of other plasmids or chromosomal DNA, or a DNA sequence integrated in the genome of an organism by CRISPR. With this platform, it would be possible to produce documentation that is clear, unambiguous, and concise.

Although individual reports for each step of a cloning strategy can be compiled using Pydnaweb, the manual nature of copy-and-paste documentation is error-prone and can result in prolonged documentation efforts. To mitigate these issues, we developed dnaudit, a tool designed to verify the completeness and accuracy of a planned cloning strategy. In addition to validation, dnaudit offers an integrated overview of how individual unit operations are connected, streamlining the documentation process and reducing the likelihood of errors.

Neither Pydnaweb nor dnaudit is strictly necessary for interpreting a cloning strategy. However, using these two stakeholder-driven services/tools in a complementary manner significantly facilitates the production of complete and accurate documentation that meets the FAIR principles (17). These tools will, in turn, ensure that the documentation remains easy to save, track, and share, with no tool lock-in, allowing for flexibility and long-term usability.

### 2.2. Specifications of Pydnaweb / dnaudit

The documentation consists of text files (snippets) that each outline a molecular biology unit operation, such as PCR, Homologous Recombination, or CRISPR digestion. These snippets should have a text “.txt” or markdown “.md” file extension. Each unit operation includes a series of input sequences in a specific order, along with at least one resulting sequence, all of which must be in FASTA format. Sequences should be identified by a unique name or identifier.

Each unit operation should begin with a title with one of the reserved words from a very reduced, easy-to-remember collection of headers (**Table 2**), optionally preceded by yaml front matter. Restriction enzyme names, such as *BamHI* or *HindIII*, should be written exactly as they appear in REBASE (http://rebase.neb.com). Additionally, reserved key/value expressions are associated with each sequence, indicating checksum, strandedness (single or double), topology (linear or circular) or alphabet (IUPAC or dsIUPAC). These are located after the identifier in the FASTA header or within the comment section of a GenBank file (**Table 3**).

**Table 2:** Keywords from the collection of headers for some of the molecular biology unit operations.

| Header | Molecular biology unit operation |
| --- | --- |
| # pcr | PCR reaction |
| # cut | Restriction digestion |
| # ligate | Ligation with a DNA ligase |
| # homologous_recombination | Homologous recombination |
| # crispr | CRISPR digestion |
| # fusion_pcr | Fusion PCR |

**Table 3:** Reserved value pairs indicating checksum, strandedness, topology or alphabet of each sequence included in each snippet.

| Reserved Value Pairs | Meaning | Note |
| --- | --- | --- |
| cdseguid=... | Checksum for a circular dsDNA sequence | Implies a circular molecule even in the absence of topology=... |
| ldseguid=... | Checksum for a linear dsDNA sequence | Implies a linear molecule even in the absence of topology=... |
| csseguid=... | Checksum for a circular ssDNA sequence | Implies a circular molecule even in the absence of topology=... |
| lsseguid=... | Checksum for a linear ssDNA sequence | Implies a linear molecule even in the absence of topology=... |
| alphabet=dsIUPAC | Genetic code is dsIUPAC | Default genetic code is IUPAC |

All relevant files should be organized within the same project folder structure to ensure consistency and easy access to the complete documentation of a molecular biology cloning strategy. Sequences are identified by their unique name or identifier, which may differ from the file name. All file names within the project folder must be unique, and each DNA sequence can only have one identifier in the project.

Pydnaweb provides a series of tools (**Table 4**), each simulating a molecular biology unit operation and producing a cloning strategy snippet directly compatible with dnaudit. Nevertheless, each tool can be used separately.

**Table 4:** Some simulation, design, and utility tools available in Pydnaweb as an online service (available in https://pydnaweb.streamlit.app/assembly).

| Simulation/Design/Utility Tool | Description |
| --- | --- |
| pcr | Simulate PCR (if a pair of primers and a template sequence are already available) |
| cut | Simulate restriction digestion with a chosen enzyme |
| crispr | Simulate CRISPR digestion with pre-designed guide RNA and known target sequence |
| assembly | Simulator for linear or circular assembly of a list of sequence fragments |
| fusion pcr | Simulate fusion PCR based on homology between two DNA templates |
| gateway | Simulate Gateway cloning |
| Primer design | Design a new pair of primers for a template sequence or a list of template sequences |
| Assembly design | Design primers for linear or circular assembly of a list of amplicons or sequence fragments |
| Design matching primer | Design a primer to match an existing primer for a primer/template pair or list of such pairs |
| Enumerate primer list | Assign numbers to primers in a list (FASTA or GenBank format); numbers will be prepended to each identifier |
| Seguid calculator | Calculate SEGUID checksums for protein or DNA sequences |
| tab format | Format primer list in tab format. Required for many companies selling oligonucleotides |
| Toggle format | Reformat sequences in GenBank or FASTA format |

### 2.3. Pydnaweb for the simulation of gene deletion in *Saccharomyces cerevisiae*

The *S. cerevisiae* phosphoglucose isomerase 1 enzyme interconverts glucose 6-phosphate and fructose 6-phosphate, therefore playing a crucial role in both the glycolysis and gluconeogenesis pathways (19). The gene encoding for this enzyme was disrupted by integrating a natMX6 cassette in the *PGI1* locus, which would confer the *pgi1*Δ strain resistance to the antibiotic nourseothricin. Briefly, the natMX6 cassette from the pAG25 plasmid (Addgene plasmid # 35121; 20) was amplified by PCR using primer *383_up-PGI1* and primer *382_dn-PGI1* (**Table S1**). Homology arms for the up and downstream regions of the *PGI1* gene were added to the ends of the PCR product by tails added to the primers. The PCR product was transformed into *S. cerevisiae* cells to disrupt the *PGI1* gene in its genome (**Figure 2**).

**Figure 2:**
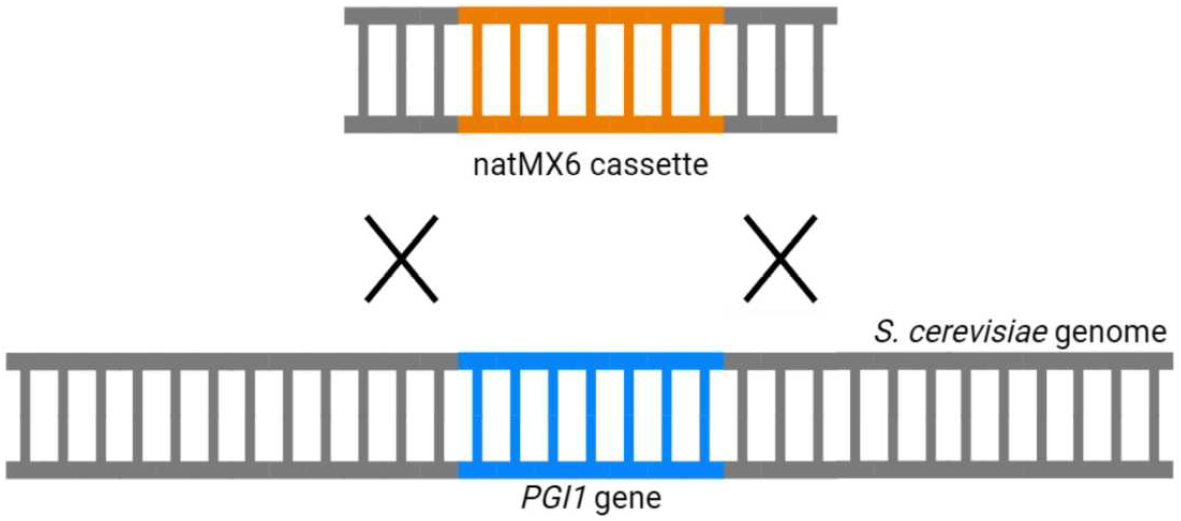
Schematic representation of gene replacement in *S. cerevisiae* via homologous recombination. The *PGI1* gene (blue) in the yeast genome is targeted for disruption by the natMX6 cassette (orange), which confers resistance to NAT. Homologous recombination at the flanking regions (indicated by Xs) facilitates the replacement of the *PGI1* gene with the natMX6 selectable marker.

The “pcr_natMX6_cassette_pAG25” snippet **(Figure 51)**, obtained from the WebPCR simulating tool from Pydnaweb (https://pydnaweb.streamlit.app/assembly). describes the PCR reaction for amplifying the natMX6 cassette. The primers *383_up-PG/1* and *382_dn-PG/1* **(Table 51)** were listed first, followed by the template {pAG25 plasmid) and the PCR product last. The order of entries is important, and a blank line must be left between sequences and any associated comments. Note that the PCR program suggested by WebPCR and the used enzyme for the elongation step were also incorporated in the snippet, so this amplification strategy is fully documented for reproducibility.

Following the amplification of the desired PCR product, homologous recombination was simulated using the “Assembly” tool, with the amplified DNA sequence and the target locus provided as inputs.

Ultimately, the success of the deletion was confirmed by cell growth in the presence of nourseothricin (NAT).

### 2.4. Pydnaweb for simulation of chromosomal GFP fusion by CRl5PR in a pathogenic fungus

Ato proteins are suggested to play a critical role in the pathogenesis of *Candida* species (21). As part of the acetate transporter (AceTr) family, they are proposed to mediate short-chain fatty acids (SCFAs) uptake (22), in particular Ato1 (23), and show strong upregulation following phagocytosis (24). Comparative phylogenetic analysis of S. *cerevisiae* Ady2/Ato1 orthologues has identified ten Ato-like proteins in C. *albicans*, indicating a significant expansion of this protein family relative to other fungal species (22).

The *C. albicans* gene *ATO1* (Accession number XP_710295.1) was fused at its 3′ end with the *GFP* gene to generate a traceable Ato1-GFP fusion protein in the *C. albicans* SC5314 strain, following an adapted version of the recyclable CRISPR-Cas9 system protocol developed by the Hernday lab (25,26). This system consists of three key components: (a) CAS9, encoding an RNA-guided endonuclease; (b) a guide RNA (gRNA) that directs CAS9 to introduce double-stranded breaks at a specific genomic locus; and (c) donor DNA (dDNA), which facilitates repair of these breaks via homologous recombination.

The CRISPR-Cas9 strategy employed in Ghasemi et al. (2025) utilizes the nourseothricin-resistance marker (NAT) to select for stable expression of Cas9 and gRNA. This method follows a split-marker approach, in which the Cas9 and gRNA expression constructs are carried on separate plasmids, each containing overlapping fragments of the NAT marker (25,26). All steps including this genetic manipulation were simulated using the Pydnaweb tools “pcr”, “fusion pcr”, “cut”, “crispr” and “assembly”.

To generate the gRNA cassette (Fragment C), two intermediate PCR fragments - A and B - must first be assembled. Fragment A (**Figure 3.A**), a universal fragment containing the 2-of-2 portion of the NAT marker, is amplified from plasmid pADH110 (Addgene plasmid # 90982) using primers AHO1096 and AHO1098 (**Table S1**). Fragment B (**Figure 3.B**), which includes the custom gRNA and the downstream region of the *HIS1* ORF, is amplified from pADH147 (Addgene plasmid # 90991) using a custom gRNA oligo and primer AHO1097 as forward and reverse primers (**Table S1**), respectively. These two fragments are then fused by overlap-extension PCR using oligonucleotides AHO1237 (C-6423, forward primer) and AHO1236 (C-6422, reverse primer) (**Table S1**), yielding the final gRNA cassette (Fragment C, **Figure 3.C**).

**Figure 3:**
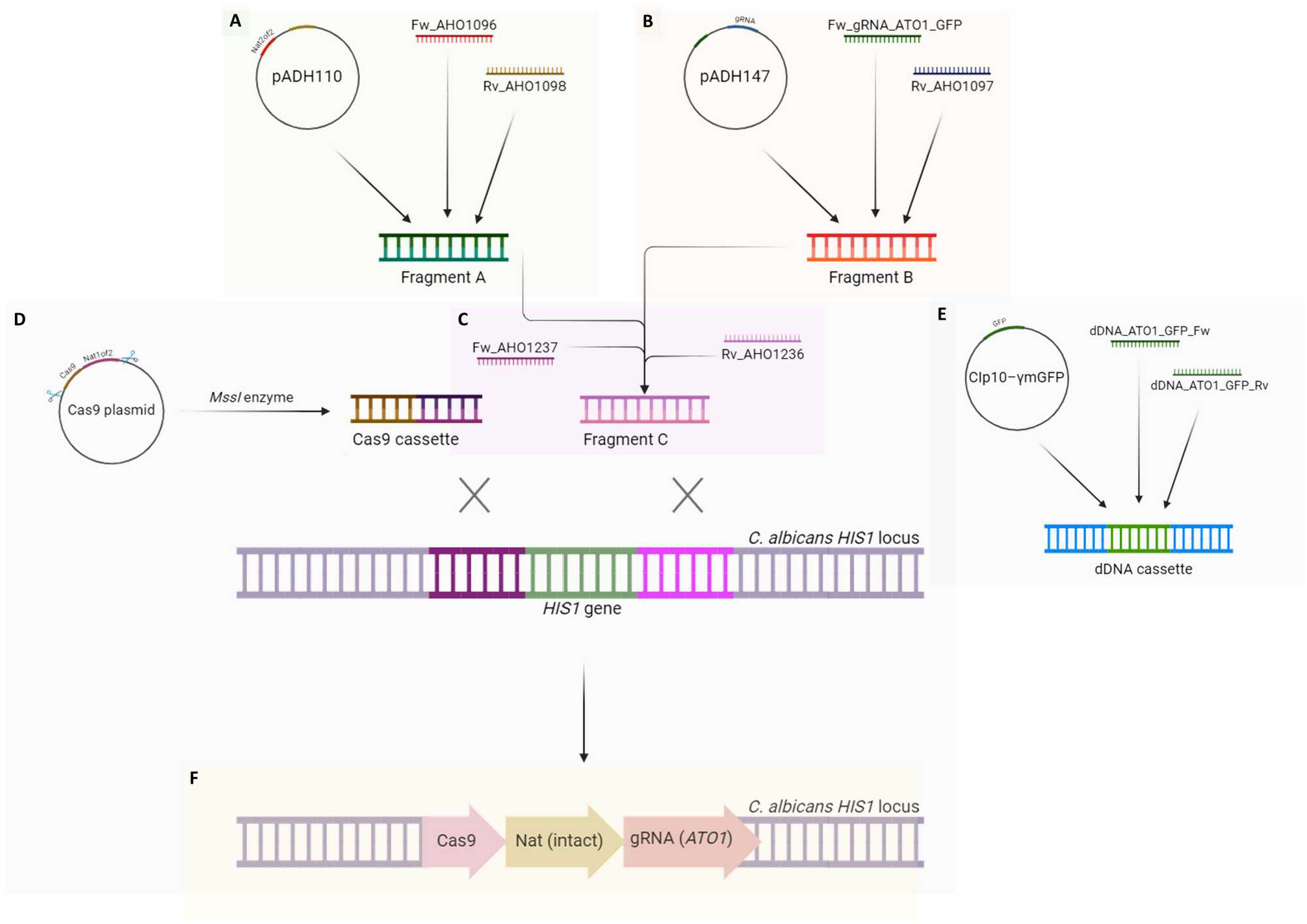
Visual representation of the steps involved in the cloning strategy for a GFP fusion in the 3’ end of the *ATO1* gene in *C. albicans* at the chromosomal level. **A**. amplification of Fragment A by PCR; **B**. amplification of Fragment B containing the gRNA by PCR; **C**. amplification of Fragment C by fusion PCR; **D**. digestion of the Cas9 plasmid using the *MssI* restriction enzyme to generate the final Cas9 cassette; **E**. amplification of the dDNA cassette (GFP) by PCR; **F**. integration of the Cas9 cassette and Fragment C in the *C. albicans HIS1* locus by homologous recombination.

Finally, Fragment C was co-transformed into *C. albicans* SC5314 together with the appropriately digested CAS9 pADH99 plasmid (with *MssI* restriction enzyme; Addgene plasmid # 90979; **Figure 3.D**) and the donor DNA. dDNAs (**Figure 3.E**) were amplified from the template CIp10–γmGFP (Addgene plasmid # 163119, 27) using the oligonucleotides *dDNA_ATO1_GFP_Fw* and *dDNA_ATO1_GFP_Rv* as forward and reverse primers, respectively (**Table S1**).

During transformation, homologous recombination between the overlapping NAT1-of-2 and NAT2-of-2 fragments restores a functional NAT resistance marker, while upstream and downstream regions of both cassettes allow for genomic integration (**Figure 3.F**). As a result, only transformants that successfully integrated the complete NAT cassette will grow on selective media.

Furthermore, the dDNA cassette containing flanking regions matching the target sequence may be integrated into the double-stranded break (DSB) site (**Figure 4**).

**Figure 4:**
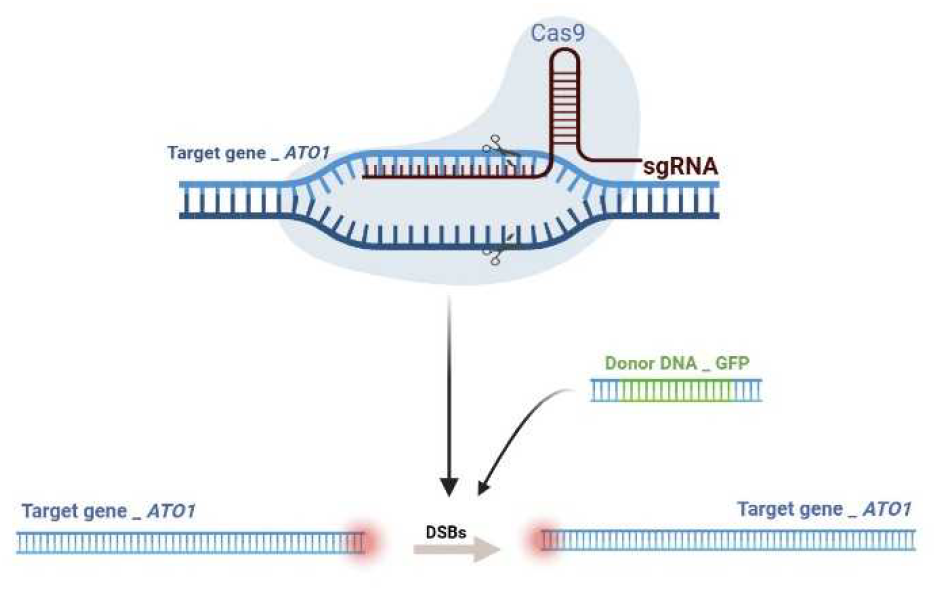
CRISPR-Cas9 mediated gene editing of the *ATO1* target gene. The Cas9 endonuclease is guided by the gRNA to the target site within the *ATO1* gene. The gRNA binds to the complementary DNA sequence, allowing Cas9 to introduce a double-strand break (DSB) at that locus (Figure S2). The donor DNA cassette containing the gene encoding GFP facilitates homology-directed repair (HDR), resulting in the insertion of the GFP sequence into the 3’ end of the *ATO1* gene.

It is important to note that the design of the dDNA did account for the possibility that, following transformation, the same gene could still be targeted by the CRISPR-Cas9 system. To address this, a simulation using the “crispr” tool was also performed to assess CRISPR targeting after the insertion of GFP downstream of the *ATO1* gene (**Figure S3**). Since no CRISPR digestion was predicted, we could assume that Cas9 activity at the modified locus was prevented. This suggests that the gene edit is stable and unlikely to be re-targeted by the CRISPR system.

To study the expression and subcellular localization of the Ato1-GFP fused protein, we examined *C. albicans* cells expressing Ato1-GFP during direct growth on 0.1 % (v/v) succinic acid, pH 5.0 (**Figure 5**). We could then confirm that the genetic construct was correctly assembled, as it enabled us to track the intracellular trafficking of the Ato1 protein under specific conditions. This validation highlights the importance of using Pydnaweb to design and simulate the cloning process. By virtually modeling each cloning step - including Restriction digestions, PCRs, and Assemblies - we were able to identify potential design flaws, verify primer-target compatibility, and predict the final sequence of the recombinant DNA prior to any wet-lab experimentation. This computational approach not only minimized trial-and-error but also enhanced the overall reliability and reproducibility of the cloning workflow.

**Figure 5:**
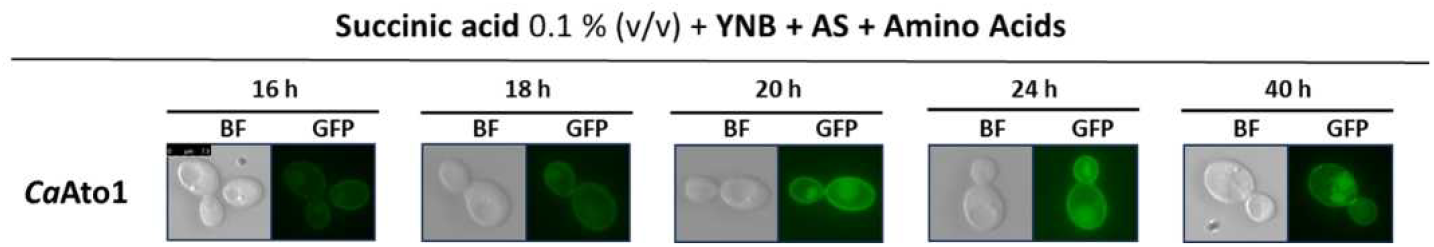
Subcellular localization of Ato1-GFP protein. *C. albicans* cells were grown directly in 20 mL SC medium supplemented with 0.1 % (v/v) succinic acid, pH 5.0, with an initial OD _640nm_ of 0.0005. When cells reached an OD_640nm_ between 0.3 and 0.6 (t ≈ 16 hours), they were visualized by fluorescence microscopy. Subsequently, samples were observed after 18, 20, 24, and 40 hours of growth. Cell cultures were grown with aeration at 200 rpm, at 30 °C. The abbreviations BF and GFP refer to “Bright-Field” and “Green Fluorescent Protein,” respectively.

### 2.5. dnaudit: Verifying reproducible cloning workflows

This section demonstrates the utility of dnaudit in verifying cloning workflows. As a case study, we used well-documented snippets that simulate a PCR using the *S. cerevisiae* Cytochrome c isoform 1 gene as the template, with primers *1_5CYC1clone* and *2_3CYC1clon* serving as the forward and reverse primers, respectively (**Table S1**). The resulting PCR product is then used as the template for a second amplification, this time with primers *f168* and *r168* (**Table S1**).

The results are visualized through a NetworkX graph (https://networkx.org/) that can be displayed through the gravis package, producing a d3 JavaScript-based interactive graph (**Figure 6**) that allows for rapid inspection and verification of the overall structure of the cloning strategy. dnaudit runs through collected molecular biology operation snippets (such as those describing PCR) to identify and flag any inaccuracies in the sequences. The resulting graph shows the interdependency of the unit operations and their relationships with initial, intermediate, and final sequences.

**Figure 6:**
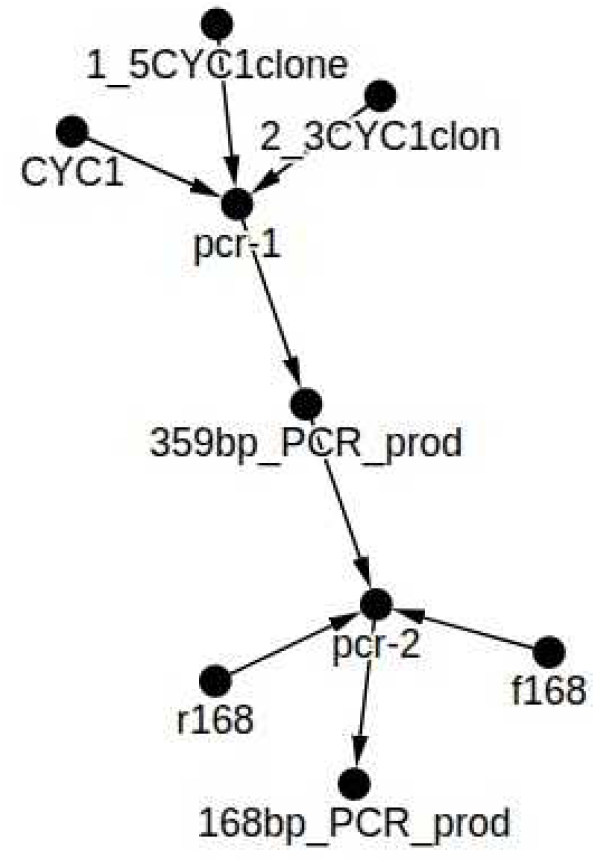
dnaudit output for verification of the overall structure of the cloning strategy. dnaudit processes FASTA sequences and/or txt./md. files and generates an interactive JavaScript-based graph. The generated graph helped in checking for inaccuracies in the cloning workflow that was designed for amplifying by PCR the *S. cerevisiae* Cytochrome c isoform 1 gene using primers *1_5CYC1clone* and *2_3CYC1clon* (Table S1). The PCR product was then used for a second PCR with primers *f168* and *r168* (Table S1).

## 3. Discussion

## 4. Acknowledgments & Funding

This work was supported by the MetaFungal project PTDC/BIA-MIC/5246/2020 (http://doi.org/10.54499/PTDC/BIA-MIC/5246/2020) through the “Fundação para a Ciência e a Tecnologia” (FCT).

Work at University of Minho (CBMA) was supported by the “Contrato-Programa” UID/04050 funded by national funds through the FCT I.P.

Work at KU Leuven (Laboratory of Molecular Cell Biology) was supported by a grant from the Fund of Scientific Research Flanders (FWO # G0C0622N).

FG acknowledges FCT for the 2023.03135.BD PhD grant (https://doi.org/10.54499/2023.03135.BD) and Erasmus+ Program for supporting her stay at KU Leuven (Belgium).

PA acknowledges FCT for the 2024.03178.BDANA PhD grant.

## 5. Competing Interest Statement

The authors declare no competing interests or financial conflicts related to the content of this study.

## 6. Supporting information

Additional supporting information is available in the Supplementary Data_1 file (PDF format).

## Notes

### Competing Interest Statement

The authors have declared no competing interest.

