## Supplementary data_1 for "dnaudit + pydnaweb: A lightweight text-based planning and documentation workflow for genetic cloning with automatic verification"

Patrícia Ataíde ^1,2^, Faezeh Ghasemi ^1,2^, Paulo César Silva ^1,2^, Cláudia Barata-Antunes ^1,2^, Sandra Paiva ^1,2^, Björn Johansson ^1,2*^

**Supplementary data 1**

Table S1

Figures S1, S2, S3

**Table S1 - List of oligonucleotides used in this work.**

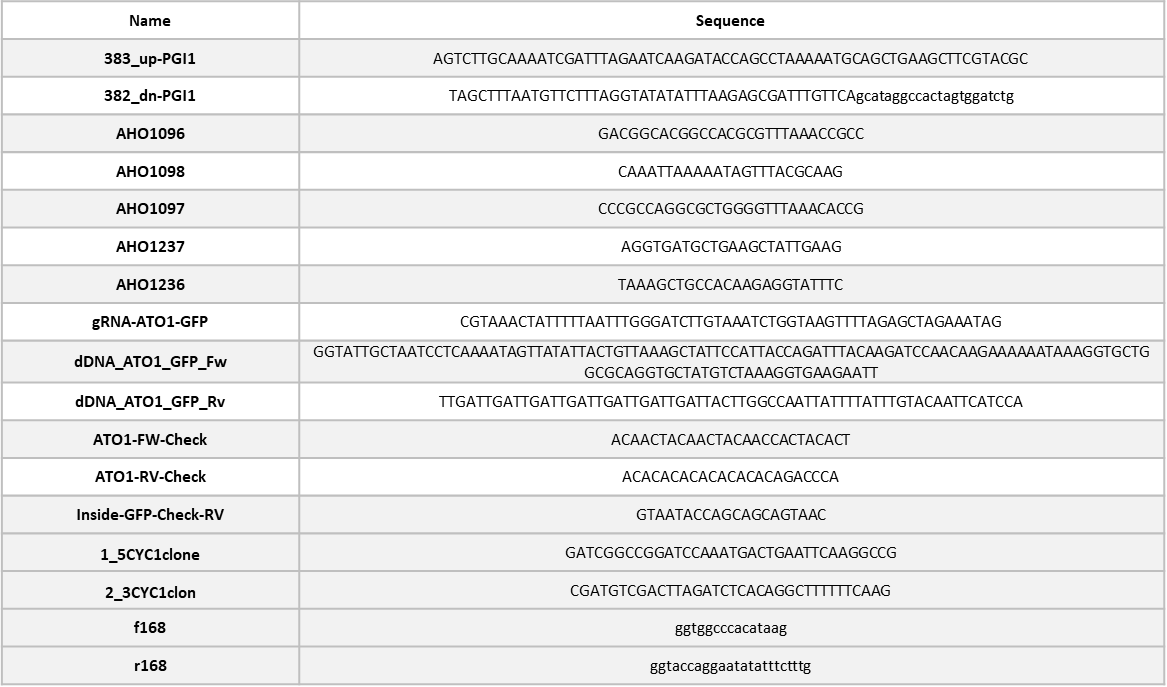

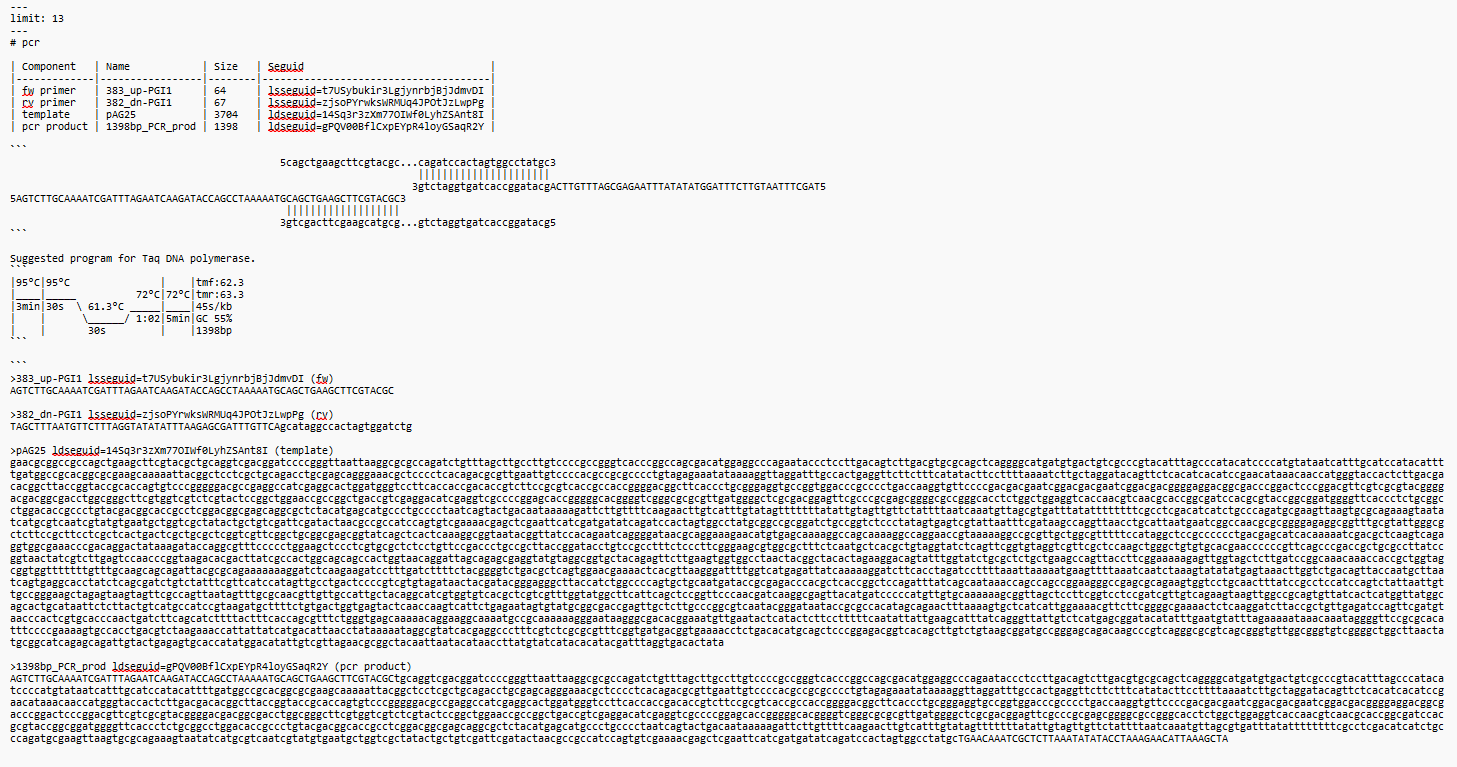

**Figure S1 - Snippet “pcr_natMX6_cassette_pAG25” obtained from the pcr simulating tool from Pydnaweb (**[**https://pydnaweb.streamlit.app/assembly**](https://pydnaweb.streamlit.app/assembly)**) describing the PCR reaction for amplifying the natMX6 cassette.** A component list identifying forward (383_up-PG11) and reverse (382_dn-PG11) primers, the template plasmid (pAG25), and the expected PCR product is present, preceded by the header “pcr”, which identifies the unit operation that will be described. Moreover, a visual alignment of the primers to the target template sequence with base-pair matching is displayed, with a suggested thermal cycling protocol for Taq DNA polymerase based on the melting temperatures (Tm) of the primers and the length of the amplified product. Lastly, full sequences of the primers, the plasmid template, and the resulting PCR product, along with their respective value expressions are also listed.

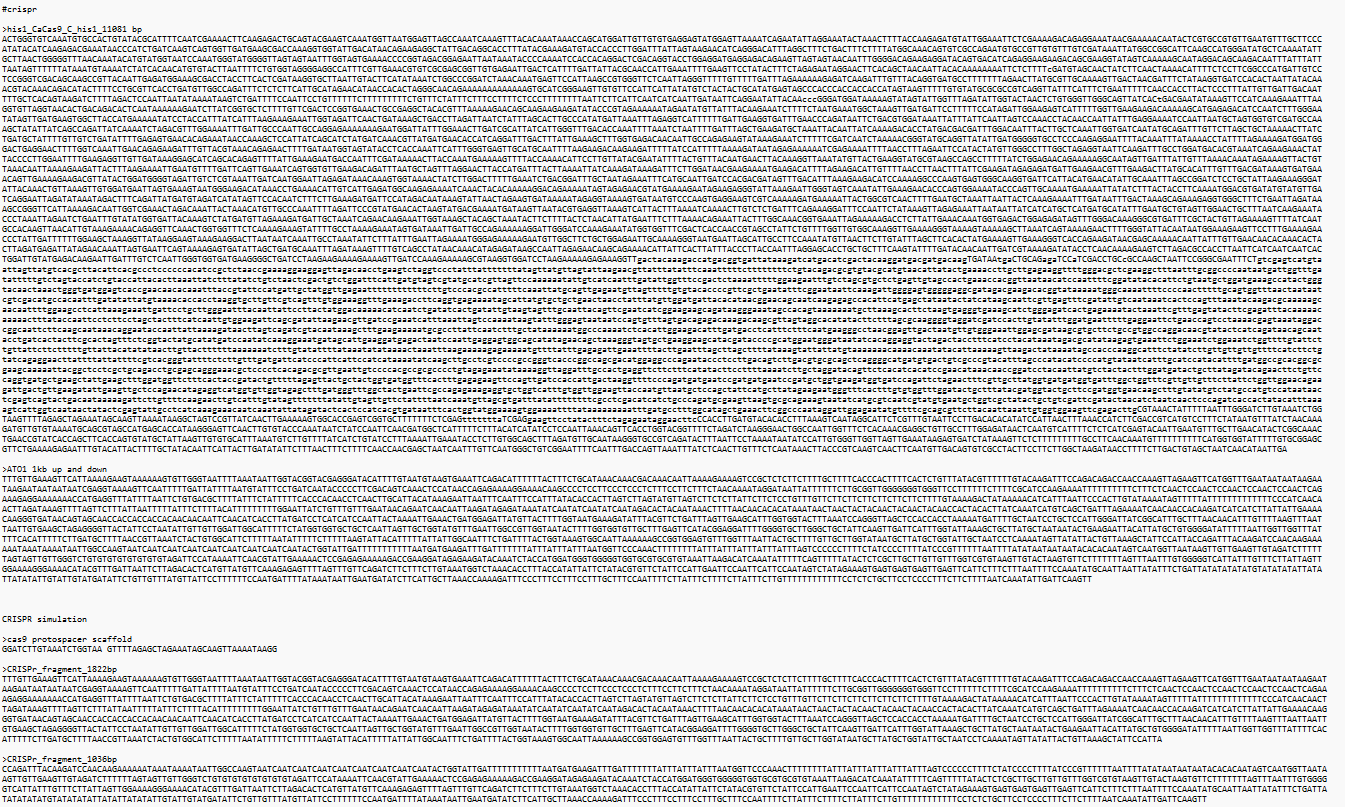

**Figure S2 - Snippet “crispr_ATO1” obtained from the crispr simulating tool from Pydnaweb (**[**https://pydnaweb.streamlit.app/assembly**](https://pydnaweb.streamlit.app/assembly)**).** As inputs, the gRNA construct and the target gene are listed first, preceded by the header “crispr”, which indicates the unit operation that will be described. The cas9 protospacer scaffold is also displayed, followed by the fragments that are generated after the creation of double-stranded breaks at the *ATO1* locus.

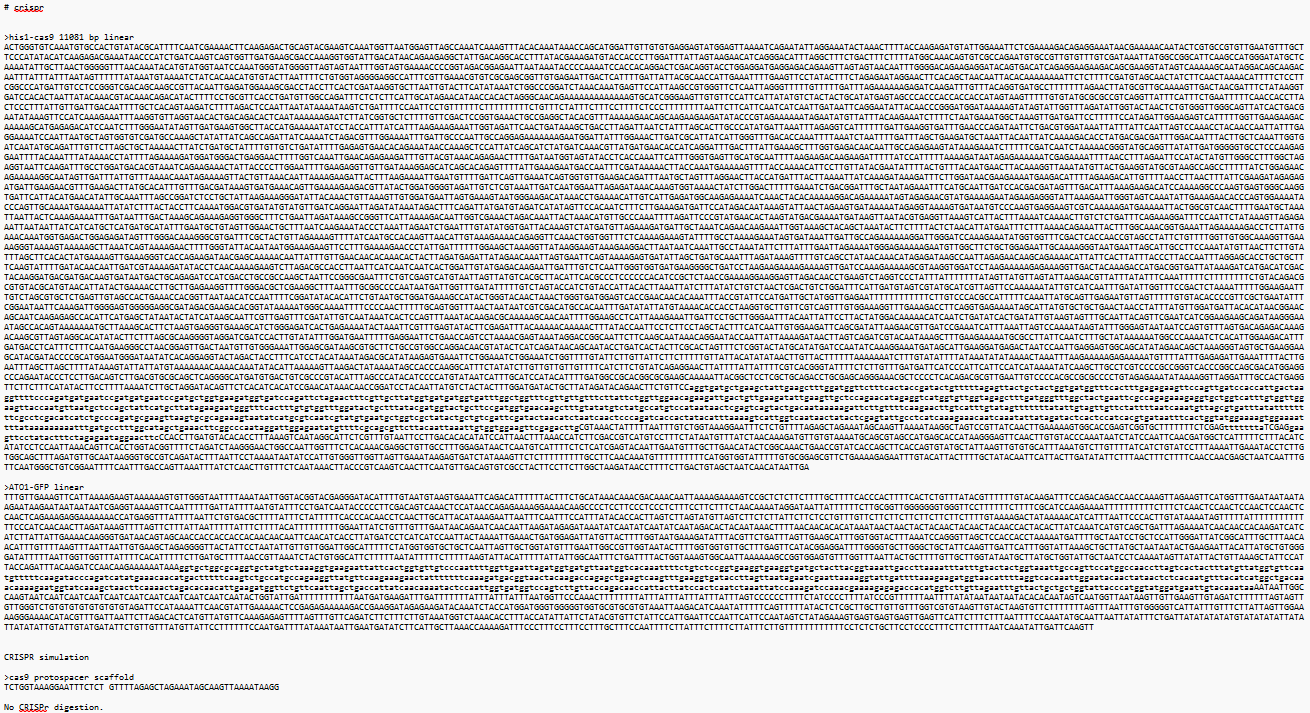

**Figure S3 - Snippet “crispr_ATO1-GFP” obtained from the crispr simulating tool from Pydnaweb (**[**https://pydnaweb.streamlit.app/assembly**](https://pydnaweb.streamlit.app/assembly)**).** As inputs, the gRNA construct and the target gene fused with the GFP gene sequence are listed first, preceded by the header “crispr”, which indicates the unit operation that will be described. The cas9 protospacer scaffold is also displayed, followed by the predicted fragments that are generated by the CRISPR system at the *ATO1* locus. No CRISPR digestion was predicted, which strengthens the hypothesis that the *ATO1* gene fused with the GFP gene sequence is no longer a target for the Cas9 enzyme after transformation and genomic integration of the dDNA (GFP).
